# LncRNA SNHG1 suppresses LPS-induced acute lung injury by regulating miR-421/TIMP3 axis

**DOI:** 10.1101/2021.02.04.429871

**Authors:** Zeyu Jiang, Jinyi Tan, Yan Yuan, Jiang Shen, Yan Chen

## Abstract

Extensive evidence has revealed the crucial roles of long non-coding RNAs (lncRNAs) in acute lung injury (ALI). This study aimed to explore the mechanism of lncRNA SNHG1 in lipopolysaccharides (LPS)-induced ALI. RT-qPCR was employed to test the levels of SNHG1, miR-421 and TIMP3 in A549 cells. Cell viability and apoptosis were assessed by CCK-8 assay and flow cytometry. ELISA assay was adopted to examine the levels of inflammatory-related cytokines, including IL-1β, IL-6 and TNF-α. The binding sequences of miR-421 and SNHG1 or TIMP3 were predicted using starBase software. Then dual-luciferase reporter and RIP assays were adopted to verify the interaction between miR-421 and SNHG1 or TIMP3. The protein level of TIMP3 was measured by western blotting. It was found that LPS stimulation downregulated SNHG1 level and SNHG1 addition decreased viability, and induced apoptosis as well as promoted inflammatory responses in LPS-treated A549 cells. SNHG1 could sponge miR-421 and SNHG1 protected A549 cells from LPS-induced injury via inhibiting miR-421. Moreover, TIMP3 was a target of miR-421. MiR-421 silence protected A549 cells against the LPS-triggered inhibition in viability, and promotion in apoptosis and inflammatory responses. SNHG1 could upregulate TIMP3 through acting as a ceRNA of miR-421 in A549 cells. Altogether, the present study elaborated that SNHG1 inhibited LPS-stimulated ALI by modulating the miR-421/TIMP3 axis.

## Introduction

Acute lung injury (ALI) is a serious inflammatory disease of the lung with a high mortality of 35-40% that are caused by a variety of factors, including trauma, blood transfusion, infection [1]. So far, there is no effective treatment for ALI [2]. Therefore, exploring the pathophysiological mechanisms of ALI has important significance for the prevention and treatment of ALI.

Long non-coding RNAs (LncRNAs) are a group of long RNA transcripts ((> 200 nts in length) without obvious protein coding ability [3, 4]. A growing amount of evidence suggested that lncRNAs act as vital roles in the pathophysiology of multiple diseases, including ALI. For example, lncRNA MALAT1 silence inhibited LPS-stimulated ALI via suppressing apoptosis by sponging miR-194-5p/FOXP2 axis [5]. LncRNA THRIL accelerated sepsis-induced ALI by targeting miR-424 and restoring ROCK2 [6]. Recent studies have manifested that aberrant expression of SNHG1 was implicated in Inflammation-related diseases. For example, lncRNA SNHG1 was decreased in osteoarthritis and SNHG1 alleviated the inflammation of IL-1β-induced osteoarthritis via the activating p38MAPK and NF-κB pathway by sponging miR-16-5p [7]. SNHG1 suppressed ox-LDL-stimulated inflammatory response and apoptosis of HUVECs in atherosclerosis by increasing GNAI2 and PCBP1 expression [8]. However, the functional of SNHG1 in lung injury is unknown. LncRNAs serve as competing endogenous RNAs (ceRNAs) and regulate the level and biological roles of miRNAs [9]. Recent studies elucidated that microRNAs (miRNAs) acted as an essential regulatory function in ALI development [10, 11]. For instance, Li et al demonstrated that down-regulated miR-let-7e suppressed LPS-triggered ALI through suppressing pulmonary inflammation by regulating SCOS1/NF-κB pathway [12]. Ge et al showed that downregulated miR-21 ameliorated LPS-induced ALI through elevating Bcl-2 expression [13]. Huang et al indicated that addition of miR-129-5p protected against LPS-induced ALI by attenuating HMGB1 [14]. These researches elaborated that miRNAs might be a potential candidate for ALI treatment. MiR-421 was demonstrated to serve as a promotor in some cancers, such as lung cancer and colorectal cancer [15, 16]. But the function of miR-421 in ALI is unclear. MiRNAs are small non-coding RNAs that modulate gene expression at posttranscriptional levels by targeting mRNAs [17, 18]. Tissue inhibitor of metalloproteinase 3 (TIMP-3) is a member of the metalloproteinase tissue inhibitor family [19]. TIMP3 was found to be downregulated in diabetic nephropathy and miR-770-5p accelerated podocyte apoptosis and inflammation by sponging TIMP3 [20]. This study investigated the mechanisms of miR-421/TIMP3 axis in ALI.

In the current study, SNHG1 expression was analyzed in LPS-stimulated A549 cells and its function in cell viability, apoptosis and inflammatory responses were explored.

## Materials and methods

### Cell culture and LPS treatment

The human lung adenocarcinoma A549 cells (ATCC, Manassas, VA) were grown in DMEM (Sigma-Aldrich) medium containing 10% FBS (Gibco, Carlsbad, USA), 100 U/ml penicillin and 100 μg/mL streptomycin at 37 °C with 5% CO_2_. A549 cells were plated in 6-well plates, and 1 μg/ml LPS (Sigma Aldrich) or control DMSO (Sigma Aldrich) was added in medium to establish injury model [21].

### Cell transfection

The short-hairpin RNA (shRNA) targeting SNHG1 (shSNHG1, 5’-GCUGCGAGG GUAGACAUCU-3’), shTIMP3 (5’-UCCCAGGACAGGCACAGGC-3’) and scramble control shRNA (5’-UAAGGCUAUGAAGAGAUAC-3’), miR-421 mimics (5’-ACUAUGUUGACACUUUUAUCCAA-3’) with its negative control (NC mimic, 5’-ACAUCUGCGUAAGAUUCGAGUCUA-3’), and miR-421 inhibitor (5’-UGAUACAACYGAAAAUAGGUU-3’) with its negative control (NC inhibitor, 5’-UAACUAAUACAUCGGAUU-3’) were obtained from GenePharma. For SNHG1 overexpression, the SNHG1 cDNA was cloned into the pcDNA3.1 vector (GenePharma). Cell transfection was conducted with Lipofectamine^®^ 2000 (Invitrogen) After transfection for 48 h, the cells were used for subsequent analysis.

### RT-qPCR assay

Total RNAs from A549 cells were extracted using TRIzol reagent (Invitrogen,). The PrimeScript RT reagent kit (Invitrogen) was employed to synthesize cDNA. Then, RT-qPCR was performed using SYBR-Green PCR Master Mix kit (Applied Biosystems) with an ABI7500 real-time qPCR system (ABI Company, Oyster Bay, NY, USA). The expression of genes was determined using 2^−ΔΔCt^ method. U6 and GAPDH were regarded as internal controls. The following primer sequences were used: SNHG1 forward, 5’-TAACCTGCTTGGCTCAAAGGG-3’ and reverse, 5’-CAGCCTGGAGTGAACACAGA-3’; miR-421 forward, 5’ CTGTCACTCGAGCTGCTGGAATG-3’ and reverse, 5’-ACCGTGTCGTGGAGTCGGCAATT-3’; TIMP3 forward, 5’-CACGTCCAAGACTTCTTCCACCACGGC-3’ and reverse, 5’-CACTTCATCCTTCGATTCTGGAACC-3’; GAPDH forward, 5’ CATGAGAAGTATGACAACAGCCT 3’ and reverse, 5’AGTCCTTCCACGATACCAAAGT-3’; U6 forward, 5’CTCGCTTCGGCAGCACA 3’ and reverse, 5’AACGCTTCACGAATTTGCGT-3’.

### CCK-8 assay

Cell viability was measured by a CCK-8 assay. Briefly, cells were seeded into a 96-well plate at a density of 5 × 10 ^3^ cells/well. After treatments, 10 μl CCK-8 solution (Dojindo Molecular Technologies, Gaithersburg, MD) was added to the culture medium of each well and cultivated for 1h at 37 °C. Subsequently, the optical density value at 450 nm was analyzed by a microplate reader (Bio-Rad).

### RNA immunoprecipitation (RIP)

RIP was performed by Magna RIP RNA binding protein immunoprecipitation kit (Millipore). A549 cells were lysed into RIP lysis buffer. Then cell lysate were cultivated with RIP buffer including anti-Ago2 (cat. no.186733; 1:1000; Abcam) and anti-IgG (cat. no.216324; 1:1000; Abcam) were incubated with magnetic beads. The purified RNAs were then subjected to RT-qPCR analysis.

### Dual-luciferase reporter assay

The wild type (wt) and mutant type (mut) reporter plasmids of lncRNA SNHG1 and TIMP3 were designed by GenePharma (Shanghai, China). NC mimics, miR-421 mimics, NC inhibitor, miR-421 inhibitor were co-transfected with SNHG1-wt, SNHG1-mut, TIMP3-wt and TIMP3-mut into A549 cells. After 24 h, luciferase activity was measured by the dual-luciferase reporter assay system (Promega, Madison, WI, USA).

### ELISA

A549 cells culture supernatant in 96-well plates was collected, and the productions of inflammatory cytokines IL-1β, IL-6 and TNF-α were measured by ELISA kits (R&D Systems, Minneapolis, MN, USA) as instructed and quantified by normalization to protein concentrations

### Western blotting

A549 cell lysates were prepared in lysis buffer with protease inhibitors. The protein concentration was quantified via the BCA kit (Thermo Fisher Scientific). The protein was resolved by SDS-PAGE and transferred onto PVDF membranes. After blocking in 5% skim milk, the membrane was incubated with anti-TIMP3 (cat. no. ab166705; 1:500; Abcam) and GAPDH (cat. no. ab8245, 1:5000 Abcam), followed by incubation with HRP-conjugated secondary antibody (cat. no. ab205718; 1:5,000; Abcam) and the protein bands were imaged using ImageJ software (Thermo Fisher Scientific).

### Animal experiment

12 BALB/c mice (7 weeks) were obtained from Slac Laboratories (Shanghai, China). The animal experiment was approved by the First People’s Hospital of Changzhou. Mice were randomly divided into 4 groups: i control group (injection of 40 mg/kg normal saline), LPS group (injection of 40 mg/kg LPS), LPS + pcDNA3.1 group (transfection with pcDNA3.1 prior to LPS treatment) and LPS + SNHG1 group (transfection with SNHG1 prior to LPS treatment). After 48 h of transfection with a microinjector, these mice were euthanized and the right lung were removed.

Right lung was rinsed briefly using PBS buffer (Sangon Biotech Co., Ltd.), and then weighed to obtain the wet weight. The right lung was baked at 60° C for 3 days to obtain the dry weight. Wet-to-dry (W/D) ratio was calculated by dividing the wet weight by the dry weight. The myeloperoxidase (MPO) activity of lung tissues was assessed by an MPO kit ((Nanjing JianCheng Bioengineering Institute) at 460 nm. The level of SNHG1 was detected using RT-qPCR. The concentration of IL-1β, IL-6 and TNF-α were evaluated using ELISA kits.

### Statistical analysis

Data are expressed as the mean ± SD and statistical analysis was performed using SPSS 21.0 software (IBM Corp.). Statistical differences between two groups were determined using Student’s t-test, and multiple comparisons between groups were assessed by one-way ANOVA. P<0.05 indicates a statistical difference.

## Results

### SNHG1 addition improves LPS-induced ALI

To explore the biological role of SNHG1 in ALI, an ALI cell model was built in A549 cells. RT-qPCR analysis implied that SNHG1 was highly expressed after SNHG1 addition (Figure 1A). Moreover, as exhibited in Figure 1B, SNHG1 level was decreased in the LPS-treated A549 cells, which was rescued by A549 supplementation in LPS-stimulated A549 cells. Moreover, CCK-8 and flow cytometry demonstrated that LPS treatment inhibited viability and promoted apoptosis of A549 cells. However, overexpression of SNHG1 accelerated viability and repressed apoptosis in LPS-induced A549 cells (Figure 1C and D). These results indicated that addition of SNHG1 protected A549 cells against LPS stimulated injury. Also, ELISA results revealed that the levels of inflammatory cytokines (IL-1β, IL-6 and TNF-α) were elevated in LPS-treated A549 cells, while addition of SNHG1 reversed these effects (Figure 1E). In sum, the above data showed that LPS-triggered ALI could be restored by the upregulation of SNHG1.

**Figure 1.**
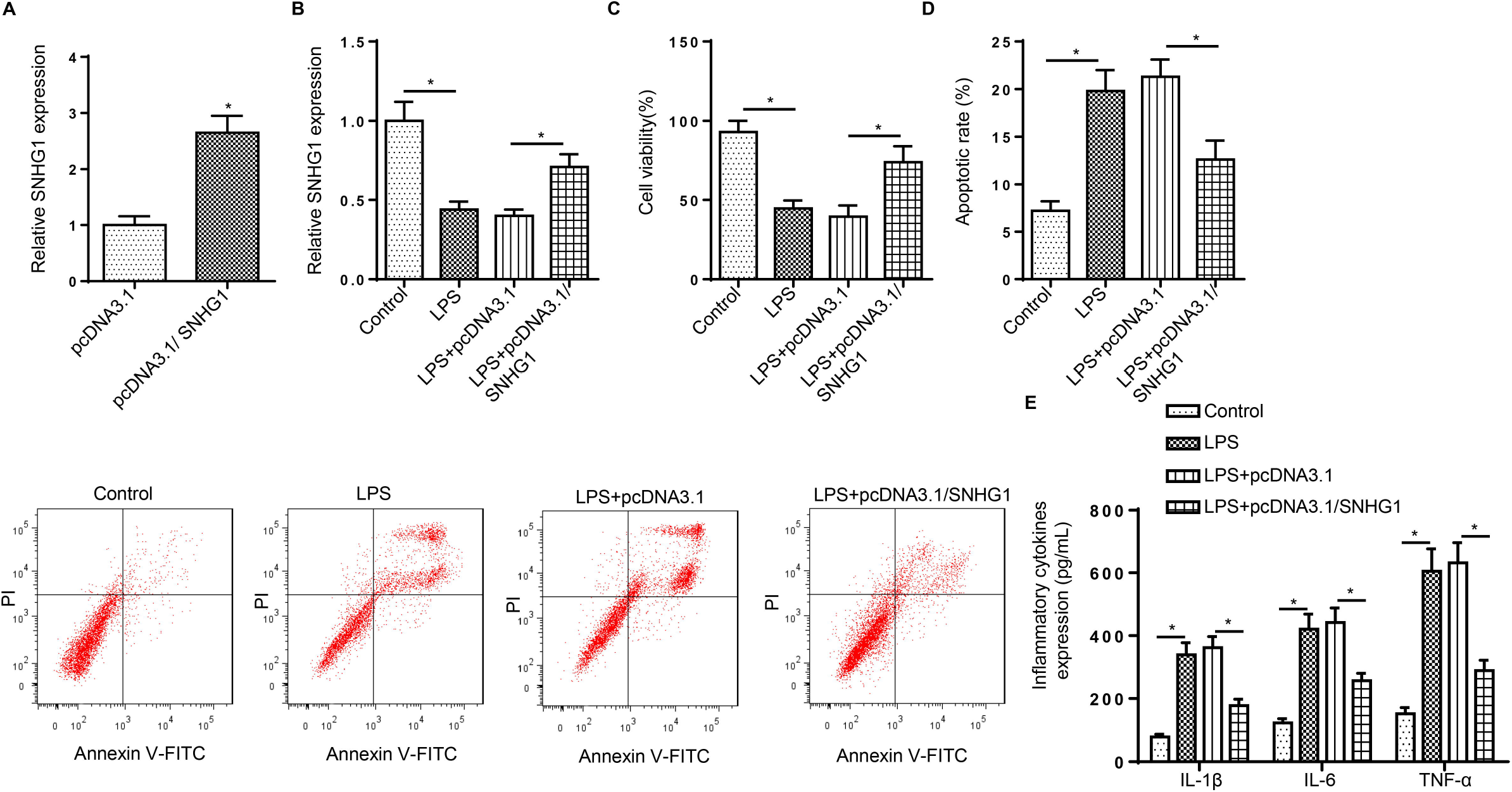
SNHG1 addition improves LPS-induced ALI. (A) RT-qPCR showed the relative SNHG1 expression in A549 cells transfected with pcDNA3.1/SNHG1. (B) RT-qPCR showed the relative SNHG1 expression in A549 cells stimulated with LPS, LPS+pcDNA3.1 or LPS+pcDNA3.1/SNHG1. (C) CCK-8 was used to determine the viability of A549 cells stimulated by LPS, LPS+pcDNA3.1 or LPS+pcDNA3.1/SNHG1. (D) Flow cytometry was used to determine the apoptosis of A549 cells stimulated by LPS, LPS+pcDNA3.1 or LPS+pcDNA3.1/SNHG1. (E) ELISA was used to determine the levels of IL-1β, IL-6 and TNF-α in A549 cells stimulated by LPS, LPS+pcDNA3.1 or LPS+pcDNA3.1/SNHG1. *p<0.05.

### SNHG1 directly bind to miR-421

StarBase software was performed to predict the potential target of SNHG1, a putative interaction between SNHG1 and miR-421 was shown in Figure 2A. Then, dual-luciferase reporter assay implied that miR-421 supplementation reduced the luciferase activity of SNHG1-wt in A549 cells, and miR-421 inhibition elevated the activity of SNHG1-wt. However, the luciferase activity in the SNHG1-mut was unaffected (Figure 2B). Moreover, RIP assay was performed to verify the interaction between SNHG1 and miR-421, and the results demonstrated that anti-Ago2 group exhibited increased levels of SNHG1 and miR-421 in A549 cells compared to anti-IgG group (Figure 2C). Additionally, RT-qPCR analysis indicated that LPS stimulation increased miR-421 level in A549 cells (Figure 2D). Deletion of SNHG1 downregulated SNHG1 in A549 cells (Figure 2E). Meanwhile, SNHG1 supplementation inhibited the level of miR-421, whereas SNHG1 depletion enhanced miR-421 expression in A549 cells (Figure 2F), indicating that miR-421 was inversely modulated by SNHG1. The data manifested that miR-421 was a target of SNHG1 and modulated by SNHG1.

**Figure 2.**
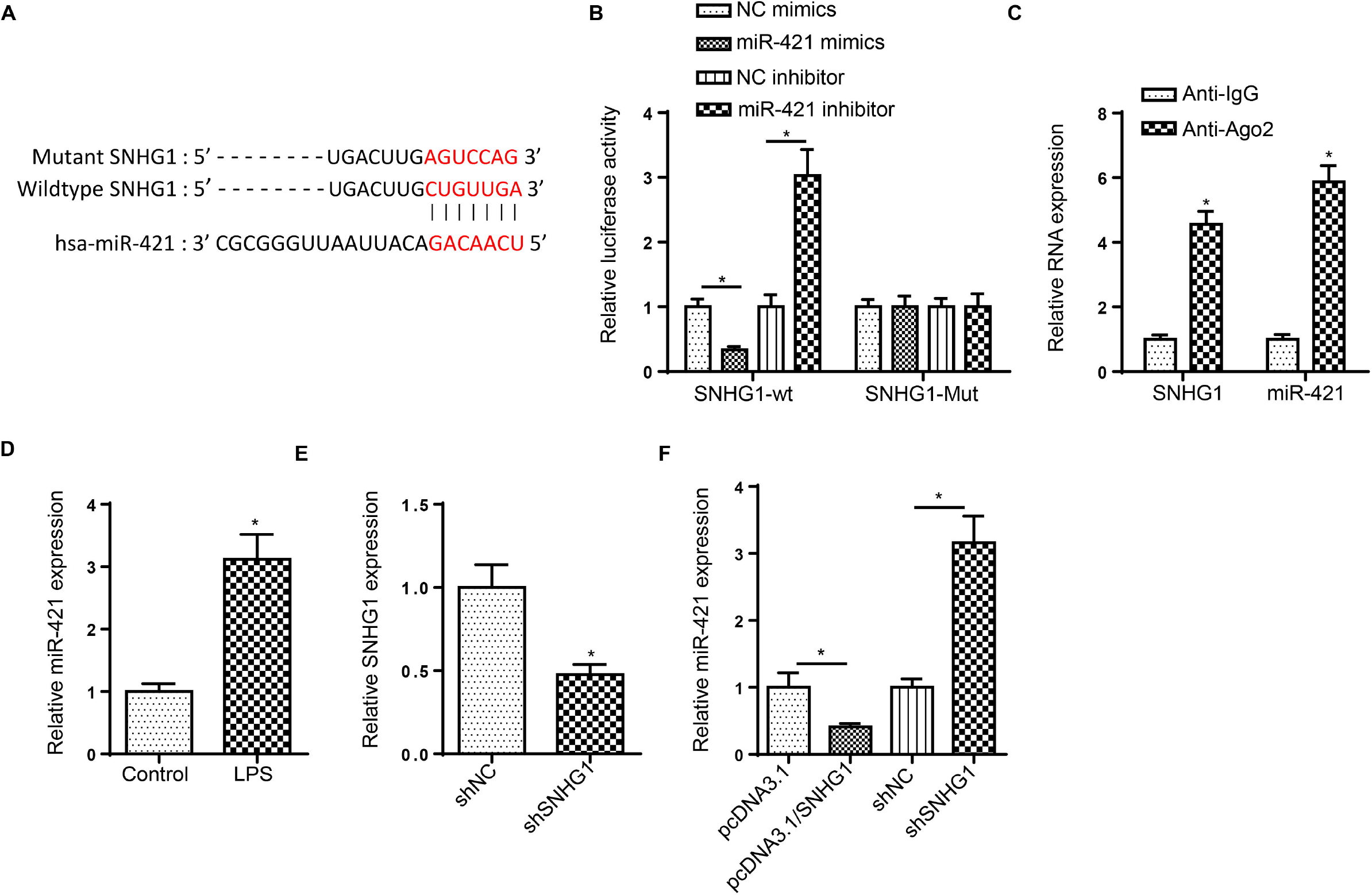
SNHG1 directly bind to miR-421. (A) StarBase website was used to predict the binding site between SNHG1 and miR-421. (B) Luciferase reporter assay showed luciferase activity of SNHG1-wt or SNHG1-mut in A549 cells transfected with NC mimics, miR-421 mimics, NC inhibitor or miR-421 inhibitor. (C) RIP assay was applied to further verify the interaction between SNHG1 and miR-421. (D) RT-qPCR showed the relative miR-421 expression in A549 cells treated with LPS. (E) RT-qPCR showed the relative SNHG1 expression in A549 cells treated with shSNHG1. (F) RT-qPCR showed the relative miR-421 expression in A549 cells treated with pcDNA3.1, pcDNA3.1/SNHG1, shNC, or shSNHG1. *p<0.05.

### SNHG1 ameliorates LPS-stimulated inflammatory damage in A549 cells by decreasing miR-421 expression

To further investigated the role of miR-421 and SNHG1 in LPS-stimulated ALI, A549 cells were treated with LPS, LPS+pcDNA3.1, LPS+pcDNA3.1/SNHG1, and LPS+ pcDNA3.1/SNHG1+miR-421 mimics. Results determined that upregulated miR-421 induced by LPS treatment was abrogated by SNHG1 addition, whereas co-transfected with pcDNA3.1/SNHG1 and miR-421 mimics elevated miR-421 level in LPS-stimulated A549 cells (Figure 3A). Moreover, addition of miR-421 abrogated the protective effect of SNHG1 supplementation on A549 cell viability (Figure 3B). Furthermore, miR-421 overexpression rescued the suppressive effect of SNHG1 supplementation on the apoptosis of A549 cells in ALI cell model (Figure 3C). Besides, we further uncovered that the repressive effect of SNHG1 addition on IL-1β, IL-6 and TNF-α levels were attenuated by miR-421 mimics (Figure 3D). Taken together, these data probed that SNHG1 repressed ALI-treated ALI via reducing miR-421 expression.

**Figure 3.**
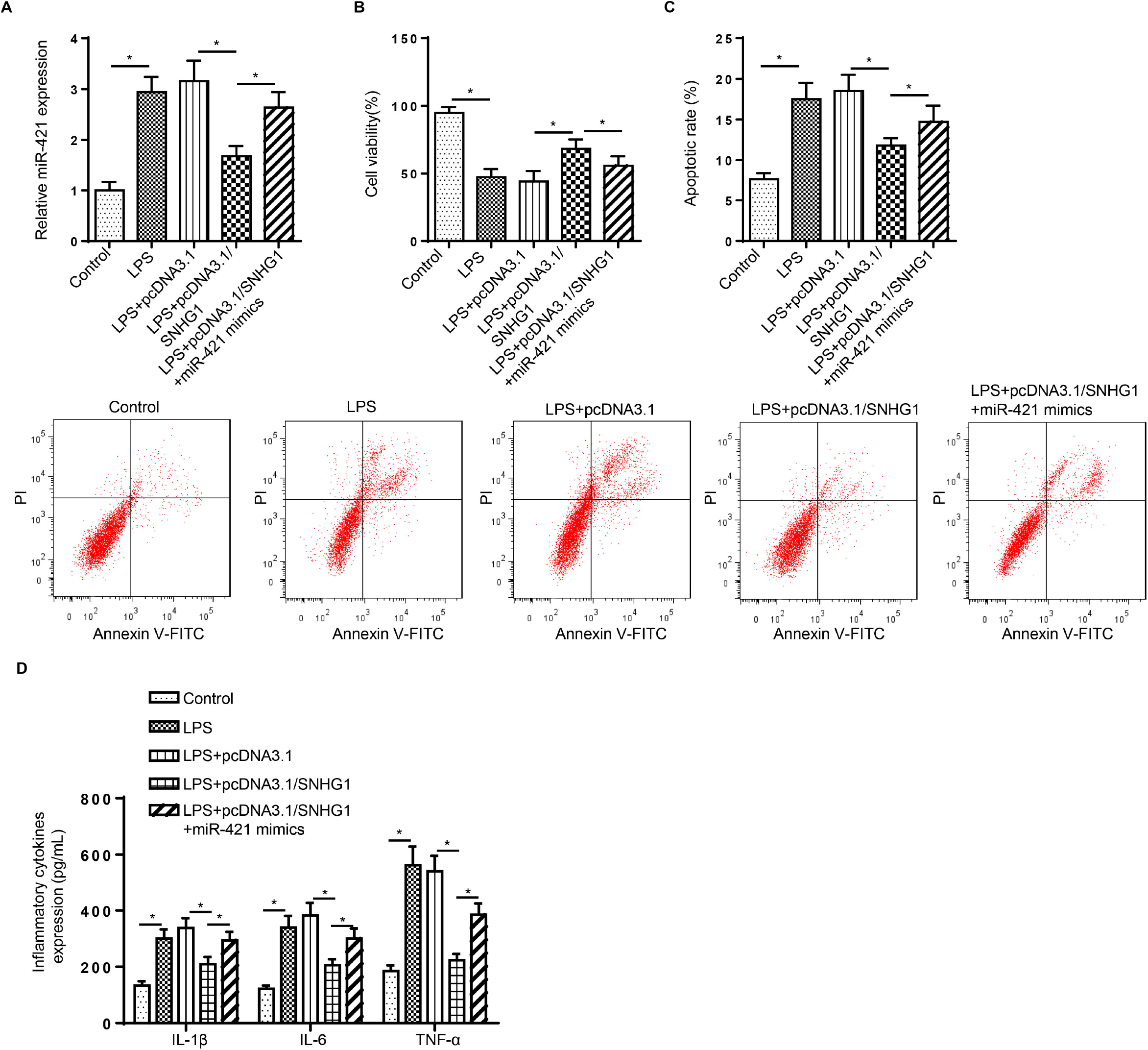
SNHG1 ameliorates LPS-stimulated inflammatory damage in A549 cells by decreasing miR-421 expression. (A) RT-qPCR was used to determine the relative miR-421 expression in A549 cells treated with LPS, LPS+pcDNA3.1, LPS+pcDNA3.1/SNHG1, LPS+pcDNA3.1/SNHG1+ miR-421 mimics. (B) CCK-8 was used to determine the viability of A549 cells stimulated by LPS, LPS+pcDNA3.1, LPS+pcDNA3.1/SNHG1, LPS+pcDNA3.1/SNHG1+ miR-421 mimics. (C) Flow cytometry was used to determine the apoptosis of A549 cells stimulated by LPS, LPS+pcDNA3.1, LPS+pcDNA3.1/SNHG1, LPS+pcDNA3.1/SNHG1 + miR-421 mimics. (D) ELISA was used to determine the levels of IL-1β, IL-6 and TNF-α in A549 cells stimulated by LPS, LPS+pcDNA3.1, LPS+pcDNA3.1/SNHG1, LPS+pcDNA3.1/SNHG1+ miR-421 mimics. *p<0.05.

### TIMP3 is a target of miR-421

StarBase software predicted that TIMP3 had a targeting relationship with miR-421 (Figure 4A). Moreover, the luciferase activity was inhibited or promoted in A549 cells co-transfected with TIMP3-wt and miR-421 mimics or miR-421 inhibitor (Figure 4B). To further validate this interaction, RIP assay was conducted. Results determined that TIMP3 and miR-421 were enriched in anti-Ago2 group in A549 cells (Figure 4C). Meanwhile, LPS stimulation decreased the mRNA and protein level of TIMP3 in A549 cells (Figure 4D). Besides, the results manifested that the mRNA and protein level of TIMP3 was inhibited by miR-421 addition, which was increased by depletion of miR-421 in A549 cells (Figure 4E). These discoveries elaborated that TIMP3 was a target of miR-421 and was negatively modulated by miR-421 in A549 cells.

**Figure 3.**
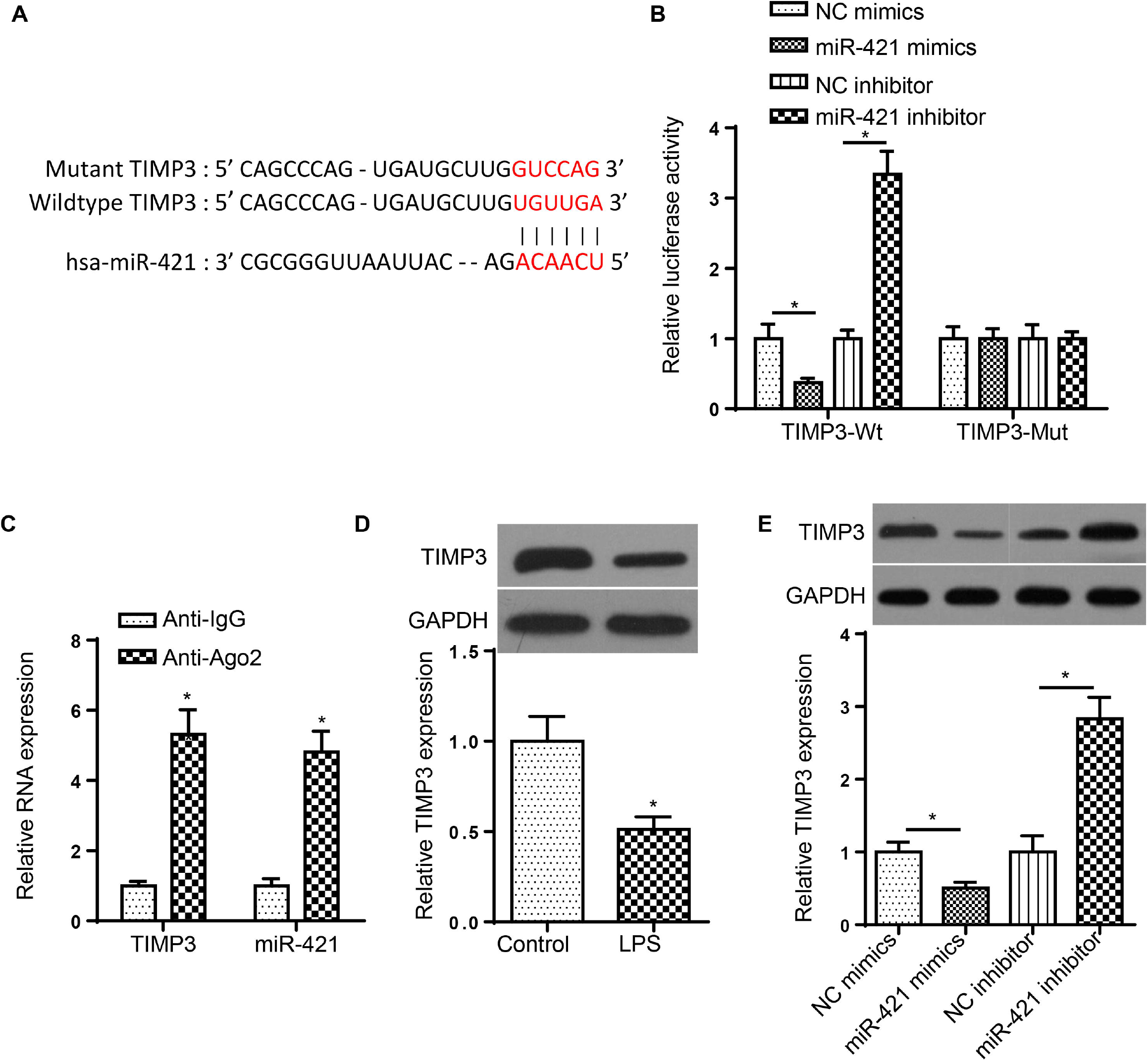
TIMP3 is a target of miR-421. (A) StarBase website was used to predict the binding site between TIMP3 and miR-421. (B) Luciferase reporter assay showed luciferase activity of TIMP3-wt or TIMP3-mut in A549 cells transfected with NC mimics, miR-421 mimics, NC inhibitor or miR-421 inhibitor. (C) RIP assay was applied to further verify the interaction between TIMP3 and miR-421. (D) RT-qPCR and western blotting assays showed the mRNA and protein expression of TIMP3 in A549 cells treated with LPS. (E) RT-qPCR and western blotting assays showed the mRNA and protein expression of TIMP3 in A549 cells treated with NC mimics, miR-421 mimics, NC inhibitor, or miR-421 inhibitor. *p<0.05.

### TIMP3 silence attenuates the protective effect of miR-421 knockdown on LPS-stimulated ALI

To probe whether TIMP3 participated in miR-421-mediated ALI, miR-421 inhibitor and shTIMP3 were co-transfected into ALI model cells. The level of TIMP3 was enhanced by miR-421 silence, and this promotion effect was abrogated by TIMP3 knockdown (Figure 5A). Moreover, miR-421 inhibition protected A549 cells against the LPS-caused the decreased cell viability and increased cell apoptosis as well as upregulated IL-1β, IL-6 and TNF-α levels. However, TIMP3 silence reversed these effects (Figure 5B-D). Collectively, miR-421 facilitated LPS-treated ALI via decreasing TIMP3 expression.

**Figure 5.**
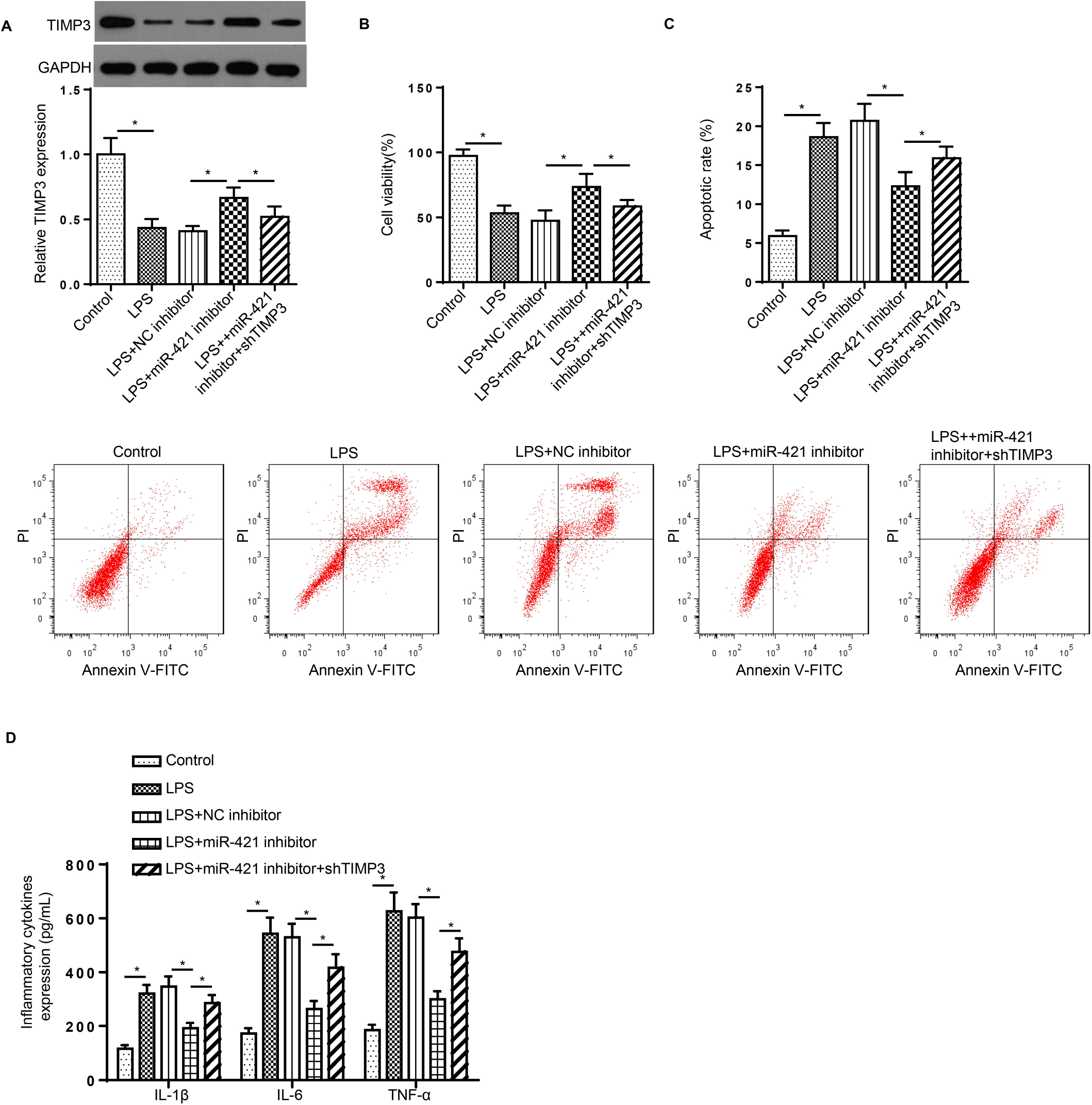
TIMP3 silence attenuates the protective effect of miR-421 knockdown on LPS-stimulated ALI. (A) RT-qPCR and western blotting assays determined the relative TIMP3 expression in A549 cells stimulated by LPS, LPS+NC inhibitor, LPS+miR-421 inhibitor, LPS+miR-421 inhibitor+shTIMP3. (B-D) CCK-8, Flow cytometry and ELISA were used to determine the viability and apoptosis as well as the levels of IL-1β, IL-6 and TNF-α of A549 cells stimulated by LPS, LPS+NC inhibitor, LPS+miR-421 inhibitor, LPS+miR-421 inhibitor+shTIMP3. *p<0.05.

### SNHG1 enhances the level of TIMP3 by sponging miR-421

RT-qPCR and western blotting assay elaborated that the level of TIMP3 was enhanced by SNHG1 addition, and miR-421 mimics decreased the level of TIMP3 (Figure 6A). Moreover, SNHG1 deficiency reduced TIMP3 expression, whereas miR-421 inhibition reversed this effect (Figure 6B). Above all, TIMP3 was modulated by the SNHG1/miR-421 axis in A549 cells.

**Figure 6.**
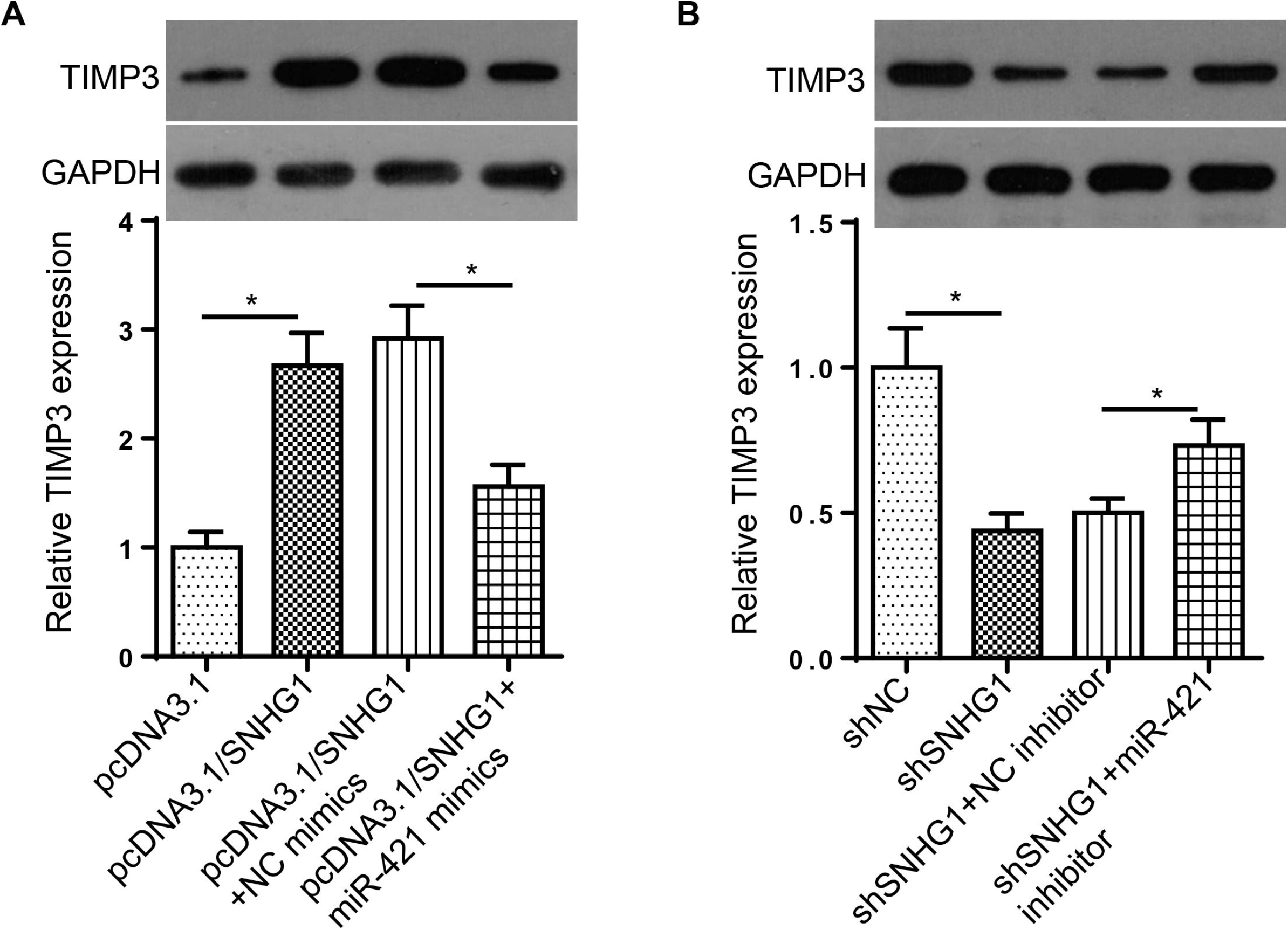
SNHG1 enhances the level of TIMP3 by sponging miR-421. (A) RT-qPCR and western blotting assay showed the relative TIMP3 expression in A549 cells transfected with pcDNA3.1, pcDNA3.1/SNHG1, pcDNA3.1/SNHG1+NC mimics or pcDNA3.1/SNHG1+miR-421 mimics. (B) RT-qPCR and western blotting assay showed the relative TIMP3 expression in A549 cells transfected with shNC, shSNHG1, shSNHG1+NC inhibitor or shSNHG1+miR-421 inhibitor, *p<0.05.

### SNHG1 addition inhibits LPS-stimulated ALI in lung tissues

Lung tissue injury was analyzed by detected the W/D ratio, MPO activity and inflammation. LPS stimulation reduced SNHG1 level, whereas supplementation of SNHG1 elevated SNHG1 expression by LPS treatment (Figure 7A). LPS stimulation increased the W/D ratio, MPO activity and inflammatory factors in lung tissues, while SNHG1 addition reversed these effects (Figure 7B-D). In summary, LPS-stimulated ALI in lung tissues was improved by the SNHG1 addition in vivo.

**Figure 7.**
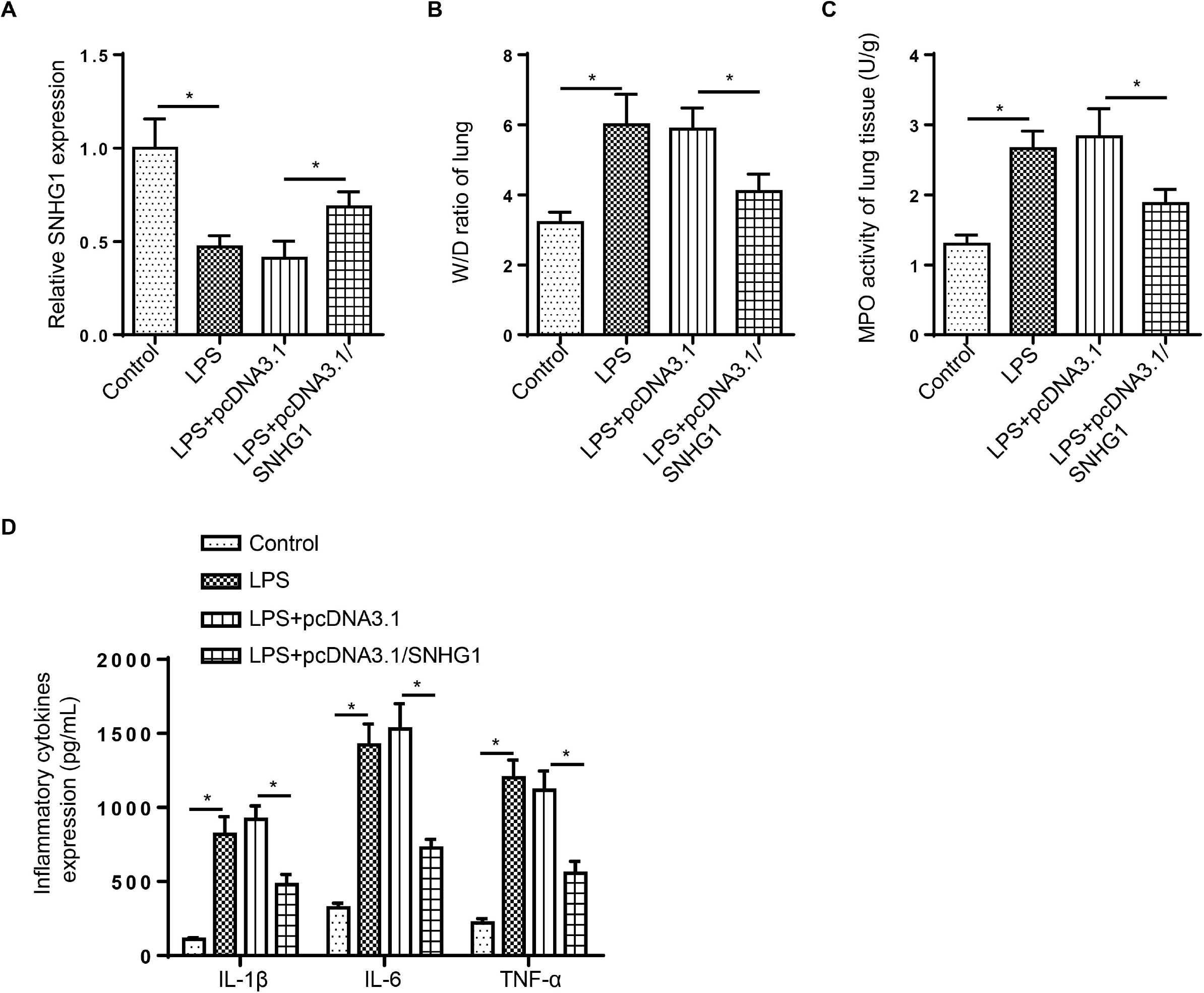
SNHG1 addition inhibits LPS-stimulated ALI in lung tissues. (A) RT-qPCR showed the relative SNHG1 expression in lung tissues of LPS, LPS+pcDNA3.1, LPS+pcDNA3.1/SNHG1 groups. (B-D) The W/D ratio and the MPO activity as well as the levels of IL-1β, IL-6 and TNF-α of lung tissues were calculated in LPS, LPS+pcDNA3.1, LPS+pcDNA3.1/SNHG1 groups. *p<0.05.

## Discussion

ALI has become a major threat to global health worldwide because of its high morbidity and mortality [22]. LncRNAs were reported to act as crucial roles in the pathophysiology of ALI. For instance, lncRNA CASC9 suppressed sepsis-induced ALI via sponging miR-195-5p and upregulating PDK4 [23]. Moreover, lncRNA TUG1 suprpressed sepsis-induced inflammation and apoptosis in ALI via regulating miR-34b-5p/GAB1 axis [24]. SNHG1 has reported to be under-expressed in ox-LDL-induced HUVECs and addition of SNHG1 suppressed ox-LDL-treated cell Injury and inflammatory response in atherosclerosis [8]. LncRNA SNHG1 was downregulated in ischemic stroke and SNHG1 addition alleviated apoptosis and inflammation via repressing miR-376a and regulating CBS/H 2 S pathway [25]. In this study, ALI cell model was constructed in.A549 cells treated by 1 μg/ml LPS and cell inflammatory response was assessed by testing the levels of IL-1β, IL-6 and TNF-α. SNHG1 inhibited cell viability and promoted apoptosis as well as increased the levels of IL-1 β, IL-6 and TNF-α in LPS-induced ALI by sponge miR-421/TIMP3 axis.

Nowadays, multiple studies have elaborated that lncRNAs can act as ceRNAs to target miRNAs to mediate the processes of various diseases [26, 27]. For example, lncRNA NEAT1 promoted the proliferation and inflammatory cytokine production by regulating miR-204-5p via NF-κB pathway in rheumatoid arthritis [28]. LncRNA LINC00240 restrained the metastasis of non-small cell lung cancer cells through sponging miR-7-5p [29]. Hence, we speculated that SNHG1 might play a regulatory role during ALI development by serving as a ceRNA. Subsequently, miR-421 was verified as a target miRNA of SNHG1. miR-421 was found to participate in the development of many types of tumors, such as ovarian cancer and papillary thyroid cancer [30, 31]. Moreover, miR-421 has been reported to aggravate inflammatory response in various diseases, such as bronchopulmonary dysplasia [32] and rheumatoid arthritis [33]. In this research, the level of miR-421 was discovered to be elevated in A549 cells by LPS treatment. Moreover, we determined that miR-421 level was inhibited by SNHG1 supplementation, which was increased by miR-421 addition. Furthermore, we further manifested that SNHG1 overexpression attenuated cell viability but increased cell apoptosis and inflammatory responses of LPS-treated A549 cells, while miR-421 addition reversed these effects. Collectively, the results suggested that SNHG1 restrained LPS-stimulated ALI through decreasing miR-421 expression.

MiRs are small non-coding RNAs that participate in physiology and disease by reducing mRNAs expression [28]. The function of miR-421 in cell biological processes are controversial because of the different target mRNAs. Therefore, we screened the possible target genes of miR-421 through starBase database and found that TIMP3 participated in multiple cellular processes, such as cell viability, apoptosis and inflammatory response [34, 35]. The current study showed that TIMP3 expression was reduced in A549 cells by LPS stimulation, and TIMP3 was inversely modulated by miR-421 in A549 cells. The acceleration of cell viability and the suppression of cell apoptosis and inflammation caused by miR-421 depletion were reversed by TIMP3 deficiency in LPS-stimulated A549 cells. Thus, these data implied that miR-421 aggravated LPS-induced ALI by downregulating TIMP3. Taken together, SNHG1 addition improved LPS-triggered ALI by regulating miR-421/TIMP3 axis.

In conclusion, this research illustrated that SNHG1 protected A549 cells against LPS-treated ALI through increasing TIMP3 expression by targeting miR-421. These results elaborated SNHG1 plays an essential role in LPS-triggered ALI and might be a novel therapeutic target in ALI.

